# Parasitoid wasp venom elevates sorbitol and alters expression of metabolic genes in human kidney cells

**DOI:** 10.1101/351031

**Authors:** Aisha L. Siebert, Luticha A. Doucette, PJ Simpson-Haidaris, John H. Werren

## Abstract

Venom from the parasitoid wasp *Nasonia vitripennis* dramatically elevates sorbitol levels in its natural fly hosts. In humans, sorbitol elevation is associated with complications of diabetes. Here we demonstrate that venom also induces this disease-relevant phenotype in human cells, and investigate possible pathways involved. Key findings are that (a) low doses of *Nasonia* venom elevate sorbitol levels in human renal mesangial cells (HRMCs) without changing glucose or fructose levels; (b) venom is a much more potent inducer of sorbitol elevation than glucose; (c) low venom doses significantly alter expression of genes involved in sterol and alcohol metabolism, transcriptional regulation, and chemical/stimulus response; (d) although venom treatment does not alter expression of the key sorbitol pathway gene aldose reductase (AR); (e) venom elevates expression of a related gene implicated in diabetes complications (AKR1C3) as well as the fructose metabolic gene (GFPT2). Although elevated sorbitol is accepted as a major contributor to secondary complications of diabetes, the molecular mechanism of sorbitol regulation and its contribution to diabetes complications are not fully understood. Our findings suggest that genes other than AR could contribute to sorbitol regulation, and more broadly illustrate the potential of parasitoid venoms for medical application.

## INTRODUCTION

Venom from the parasitoid wasp ***Nasonia vitripennis*** causes significant sorbitol accumulation in insect hosts without altering glucose level (1). Few studies have examined the conservation of parasitoid venom phenotypes in a human disease relevant model (2). Elevated sorbitol is believed to be a major contributing factor to secondary complications of diabetes (3), particularly kidney disease, which is a leading cause of morbidity and mortality in diabetics (4). However, the mechanism of sorbitol pathway regulation in the human kidney and relative contribution to glucose-induced damage is not well understood. Here we demonstrate conservation of this significant *Nasonia* venom phenotype in human cells and present a novel approach to study effects of sorbitol elevation *in vitro*, independent of glucose elevation.

Parasitoid wasps are small, abundant, and diverse hymenopteran insects with an estimated 100-400 thousand species (5–7). Female wasps inject venom into a fly host to manipulate its physiology in ways conducive to the growth of the wasp’s offspring, turning the host into an incubator with favorable biochemical composition (1, 2, 8–10). Because parasitoid wasps exploit a diverse and evolutionary divergent set of invertebrate hosts (11), it is likely that the specific venom targets within these hosts are evolutionarily conserved. There is an interest in utilizing parasitoid venoms for pharmacological applications in human disease (12, 13), although it has not been extensively explored as of yet.

Our study organism, *N. vitripennis*, is a parasitoid wasp often used as an experimental and genomic model organism (14, 15). A complete *N. vitripennis* genome (16), and partially characterized venom proteome are both publicly available (12, 17–19). Adult *Nasonia* females produce at least 93 venom peptides, including 23 novel proteins that demonstrate no homology to described sequences in other non-parasitoid organisms (17, 20). Upon injection into the fly host (*Sarcophaga bullata*), *Nasonia* venom causes several discrete metabolic changes, including a striking >20-fold increase in sorbitol with no effect on glucose levels (1). This sorbitol accumulation is preceded by induction of a subfamily of **aldo-keto reductase (AKR)** genes in the fly host, that belong to a larger family of AKRs which includes the human aldose reductase (9).

**Aldose Reductase (AR)** is annotated as the canonical sorbitol-synthesizing enzyme, it is known to convert glucose to sorbitol when glucose levels are high and it is generally assumed that elevation of sorbitol is primarily or exclusively caused by this enzyme (21–23). However, expression of a related AKR1C3 has been found to be specifically elevated in the glomerulus of patients with diabetic nephropathy (24, 25), suggesting that related enzymes may also play a role in this disease process. **Sorbitol dehydrogenase (SD)** is another important enzyme in the pathway that converts sorbitol to fructose. Insulin-insensitive cells – such as those in the kidney glomerulus, retina, and Schwann cells – express relatively little SD. As a result, these cells accumulate sorbitol when glucose to sorbitol conversion is increased (26). Inhibiting sorbitol synthesis has been a major target of drug design to prevent damage to glomerular cells (21, 23, 27–29). The relative contribution of sorbitol to this disease process is not well understood as few studies have examined sorbitol pathway regulation in human cells under biologically relevant levels of glucose (22), or independent of other potential consequences of hyperglycemia.

In this study, we investigate whether venom from the parasitoid wasp *N. vitripennis*, which dramatically elevates sorbitol in flies without altering glucose levels, can also induce elevation of sorbitol in primary **human renal mesangial cells (HRMCs)** from the glomerulus. We then compare this induction to high glucose treatment alone to assess relative efficacy of venom and glucose to elevate sorbitol. We also investigate whether venom induces targeted gene expression changes prior to sorbitol accumulation as a method of identifying potentially causative genes. Finally, we use computational methods to determine the relative binding affinity for glucose of the canonical (AR) and other candidate AKR enzymes. This study reveals a conserved venom function (sorbitol elevation) across vastly divergent species (*S. bullata* and *H. sapiens*) and provides a first step to evaluating additional candidate genes for regulating sorbitol metabolism in human cells.

## MATERIAL AND METHODS

### 2.1 Cell culture system

Human renal mesangial cells (HRMCs) were chosen because of their high rate of metabolic activity, as well as their important roles in regulating blood flow in the kidney and in the pathophysiology of diabetic nephropathy (30, 31). Within the kidney, these cells are located in close apposition to the glomerular arterioles and therefore are exposed to circulating levels of glucose in the blood. Primary HRMCs (designated P1 in our lab) were purchased from ScienCell and cultured according to the published protocol (30–32). Cells were expanded in culture to P3 and then stored in liquid nitrogen for all subsequent experiments. All **RNA-Sequencing (RNA-Seq)** and metabolic experiments were conducted on HRMCs at low passage number (<P5).

### 2.2 Wasp rearing and venom

*N. vitripennis* were reared as previously described (33, 34). Venom was collected by micro-dissection of the venom reservoir of adult female *Nasonia vitripennis* wasps per the previously published protocol (1, 18). Reservoirs were then lysed by centrifugation; venom was suspended at a concentration of 1 **venom reservoir equivalent (VRE)** per 1 ml of isotonic PBS solution and filtered through 0.22 mm PVDF centrifugal filters [Millipore-Sigma, Dramstadt Germany] to remove bacterial contamination. To produce different concentrations of venom, serial dilutions in PBS were performed. A total volume of 128 μl for each solution was added to 1 mL cell culture medium to produce different venom doses of 0, 1/128, 1/64, 1/16, 1/4, 1, 4, 16, 64, and 128 VRE per well and applied to cells grown to 70% confluence in a 24-well culture plate. Application of venom doses was randomized and all doses were applied over a five-minute period. Previous estimates of protein concentration indicate that one VRE contains approximately 1.54 μg of dry weight venom protein (35), giving a dose range of 1.2 × 10^−2^ to 197.1 mg/mL across our treatment conditions.

### 2.3 Effects of venom on HRMCs

The effect of venom was initially assessed by microscopy to examine cell count and gross phenotypic changes. Cells were photographed at baseline and again after 1 hour, 4 hours, and 24 hours of incubation with venom. Photos were trimmed, and cell count was assessed using ImageJ Analyze Particles function (36). Intracellular sorbitol level was measured, and RNA-Seq of total RNA was used to assess transcription at the same time points in samples that were plated and dosed concurrently.

### 2.4 Effects of glucose on HRMCs

To compare the effect of glucose versus venom on sorbitol level, HRMCs were cultured with escalating doses of glucose (5.5, 11, 25, 55, 110, 220 mM) dissolved in the cell culture media. Standard culture media contains 5.5 mM (1 g/L) of glucose. Glucose dose range was selected based on published work examining stimulation of the sorbitol pathway by hyperglycemia alone (37). Of note, previous experiments designed to assess activation of the sorbitol pathway routinely use the highest glucose dose (220 mM), which is equivalent to a blood glucose level of 3960 mg/dL, well above the physiological range observed in diabetic patients even during acute ketoacidosis (127-1000 mg/dL). We included these doses in order to evaluate the effects of venom relative to standard methods for sorbitol activation.

### 2.5 Sorbitol, Glucose and Fructose assays

We assessed intracellular levels of sorbitol pathway sugars by removing excess media, washing cells with isotonic PBS solution, and then lysing cells in collection solution. Assays were performed as per protocols for the colorimetric D-Sorbitol Assay Kit and Glucose Assay Kit [Adcam, Cambridge MA] to measure fructose, sorbitol, and glucose. Epoch microplate spectrophotometer [BioTek, Winooski VT]. Sorbitol elevation is then calculated relative to the sorbitol level detected in control samples (5.5 mM glucose, no venom), and similarly for fructose and glucose. To establish comparable measures of sorbitol, fold sorbitol elevation was calculated as the increase in sorbitol observed in experimental samples relative to controls pooled across all sorbitol assays (venom and glucose treatment). Protein concentration within cell lysate was used as a proxy measure of total cell density and was used to normalize sorbitol level.

### 2.6 RNA-Seq and gene expression

RNA was collected with TRIzol^®^ reagent as per manufacturer’s published protocol [Invitrogen Corporation, Carlsbad CA]. Quality measure was obtained using an Agilent 2100 bioanalyzer and all samples were determined to have an RNA Integrity of >9.0 prior to sequencing. mRNA abundance was quantified using high throughput RNA-Seq performed on an Illumina Hi-Seq2500 machine (50 base, single-end reads) with 20-25 million reads per sample, as per our previously published workflow (9, 18). The TruSeq RNA Sample Preparation Kit V2 [Illumina, San Diego CA] was used for next generation sequencing library construction per manufacturer’s protocols. The University of Rochester Functional Genomics Center performed the library preparation and sequencing according to standard Illumina protocols.

For **NGS data processing,** raw 50 bp reads were de-multiplexed using configure bcl2fastq.pl version 1.8.3. Low complexity reads and vector contamination were removed using sequence cleaner (“seqclean”) and the NCBI univec database, respectively. The FASTX toolkit (fastq_quality_trimmer) was used to remove bases with quality scores below Q=13 from the end of each read. Sequence data are available in the NCBI Sequence Read Archive (SRA) SUB4119836, BioProject ID PRJNA475146. Cleaned reads were aligned to the human genome assembly version 19/GRCh37 (38) using the Burrows-Wheeler Aligner with the default settings (39). Read counts were generated with HTSeq (40). In a pilot study, we sequenced a single replicate of all venom doses at three time points after exposure (1, 4, and 24 hours). For sorbitol elevating doses (1/128, 1/64, and 1/16 VRE) we sequenced two additional biological replicates (for a total of three) 4 hours after exposure and assessed gene expression as compared to time-matched, pooled, replicated controls (n=5). Statistical tests of **differential expression (DE)** were performed using the edgeR module in R (41–43). Design matrix was programmed to control for batch effect “model.matrix(~batch+group)”. For edgeR, significant DE cutoff was set at the p<0.05 and FDR<0.01 level as per the recommended workflow (44).

### 2.7 *In silico* protein structural prediction

AKR proteins from *Homo sapiens* were input into Phyre2 structural prediction software (45) and structures were estimated using the Intensive setting. Previously published structures of control (AKR1A1), candidate (AKR1C*, AKR1D1) and known (AR = AKR1B1, SD) *H. sapiens* sorbitol pathway enzymes were downloaded from NCBI and structures were compared to predicted structures for accuracy. Molecular configuration of ligands and cofactors were downloaded from the Research Collaboratory for Structural Bioinformatics, Protein Data Bank Ligand Expo database (46, 47). Docking potential to sorbitol pathway sugars (glucose, sorbitol, fructose) was assessed for known *H. sapiens* enzymes with iGemDock (48, 49). To assess relative docking potential of AKR enzymes, the docking algorithm was run with a population size of 200, with 70 generations, and simultaneously estimated docking of either cyclical glucose with NADPH cofactor or sucrose with NAPD+ cofactor. iGemDock predictions were generated twice: first with default settings, and second with predicted active sites from Phyre2 included. Optimal program parameters were then selected to generate all estimates of relative binding affinity.

## RESULTS

### High venom doses are associated with growth inhibition and cell toxicity

We first assessed the effects of different venom doses on HRMC growth and morphology, standardizing treatments to **venom reservoir equivalents (VRE)**. Our group and others have previously demonstrated that 1 VRE (~1.54 μg dry weight venom protein) per host is sufficient to recapitulate the developmental arrest and delayed host mortality phenotype observed in envenomated natural fly hosts, as well as to induce significant gene expression changes in conserved metabolic pathways in fly hosts (18, 35). In our initial assessment we examined a wide dose range from 1/128 up to 128 VREs, applied to cells at 70% confluence and then incubated for 1 hour, 4 hours, and 24 hours. All venom doses cause an initial decline in cell count at 1 hour, with variable recovery in growth rate (Figure 1). Sorbitol elevating venom doses (1/128, 1/64, and 1/16 VRE) resulted in an accelerated growth rate in the 4-24 hour period over controls. Signs of apoptosis including cell membrane blebbing were evident at the highest doses (64 and 128 VRE) (Figure 1). Qualitative signs of less overt cell toxicity including increased nuclear inclusions were present at somewhat lower doses (4 and 16 VRE). We observed a marked decrease in cell count at 24 hours after exposure to the highest venom doses of 16, 64, and 128 VRE; Intermediate dose 4 VRE caused a slight decrease in total cell count (Figure 1). These findings are consistent with previous observations that *Nasonia* venom causes apoptosis only at very high concentrations (2). To avoid complications of cell pathology at the high doses, we concentrated our next analyses on lower venom doses (1/128 to 1 VRE).

**Figure 1.**
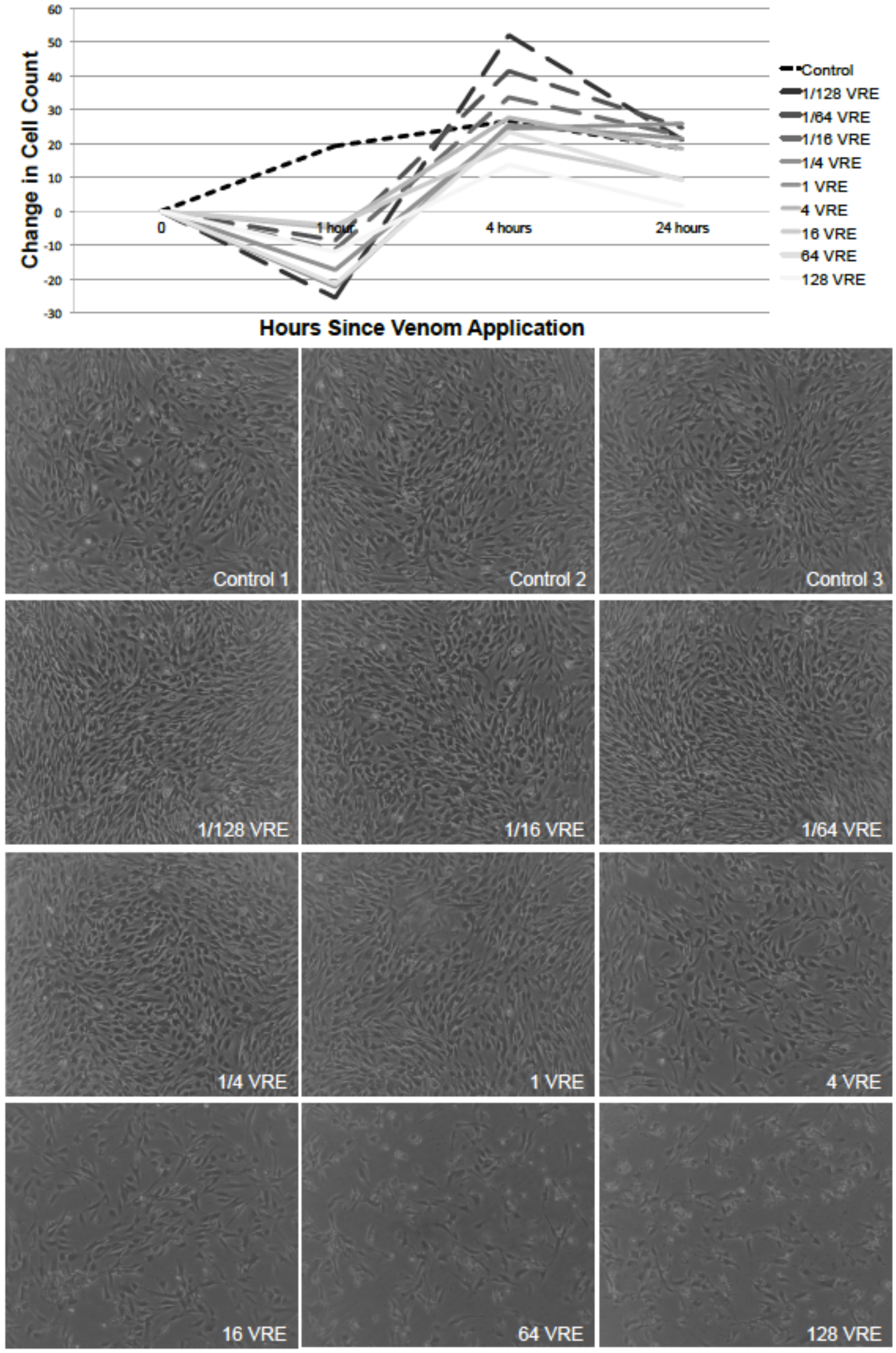
All venom doses are associated with a decrease in cell count at 1 hour, and high venom doses are associated with growth inhibition and cell toxicity at 24 hours. All venom doses cause an initial decline in cell count at 1 hour. Sorbitol elevating venom doses (1/128,1/64, and 1/16 VRE - dashed line) resulted in an accelerated growth rate in the 4-24 hour period over controls. Decreased cell count, as well as signs of apoptosis including cell membrane blebbing were evident at the highest doses (64 and 128 VRE).

### *Nasonia* venom elevates intracellular sorbitol in primary HRMCs without changing intracellular glucose or fructose levels

We assessed intracellular levels of sorbitol pathway sugars as per our experimental protocol (see Methods). After 1 and 4 hours of incubation with venom we observed no elevation of sorbitol or fructose level over that of controls for any of the venom doses (1/128 – 1 VRE). Sorbitol was markedly elevated after 24-hours of exposure, when venom caused a 5 to 25-fold increase in intracellular sorbitol levels (Figure 2). Elevation of sorbitol was greatest at the lowest venom dose (1/128 VRE - 9.6 × 10^−2^ mg per mL). Significant increases were found at 1/128, 1/16, 1/4, and 1 VRE (n=6 per venom dose; 2-tailed student’s t-test p<0.001 for 1/128 and 1 VRE, p<0.05 for 1/16 and ¼ VRE) and a non-significant increase at 1/64 (p=0.12). All venom doses 1/128 to 1 VRE significantly increased sorbitol over the controls (ANOVA p<0.0001). These results demonstrate conservation of the prominent *Nasonia* venom phenotype, sorbitol elevation, between the natural fly host *S. bullata* (1) and cultured cells from *Homo sapiens*. This phenotype coincides with an apparent absence of cell growth inhibition or visible signs of apoptosis. The finding suggests that the mechanism of venom-induced sorbitol accumulation is likely conserved, and therefore represents an important potential target for understanding sorbitol pathway regulation.

**Figure 2.**
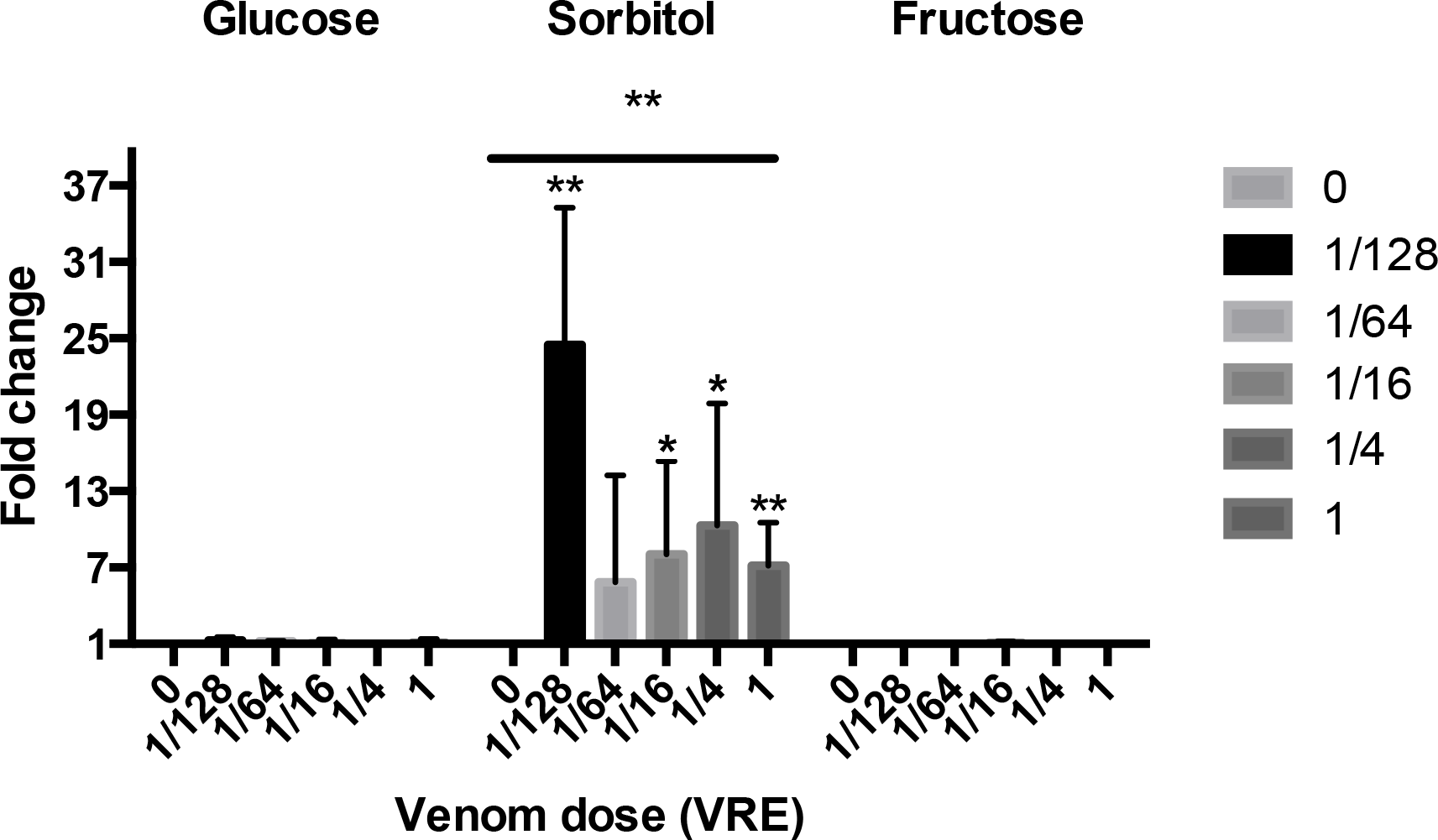
Intracellular sorbitol elevation with venom treatment. Low dose Nasonia venom elevates sorbitol but not glucose or fructose in cultured Human Renal Mesangial Cells (HRMSc) at 24 hours (ANOVA** p<0.0001; 2-tailed student’s t-test p<0.001 for 1/128 and 1 VRE, p<0.05 for 1/16 and ¼ VRE, p=0.12 for 1/64 VRE) (n=6 samples per group). Fold changes are shown relative to the zero venom control.

### Venom is a strong inducer of sorbitol elevation compared to glucose

To compared the venom phenotype to glucose-mediated sorbitol elevation, we also incubated HRMCs with escalating glucose concentrations and then assayed intracellular sorbitol levels at 24 hours. There is a significant elevation of sorbitol at the highest glucose concentrations of 110 mM and 220 mM (19.8 – 39.6 g/L) as compared to controls in standard culture media (n=4 per glucose concentration; 2-tailed student’s t-test p<0.05 for 110 and 220 mM, and not significant for 11, 25, and 55 mM) (Figure 3). Yet, peak glucose-induced sorbitol elevation was less than half that observed with low doses of venom – 1.2 × 10^−8^ g of venom (1/128 VRE) had 2.1 times the effect of 3.96 × 10^−2^ g (220 mM) of glucose on sorbitol level. **This gives venom a per unit weight efficacy in inducing sorbitol elevation that is 2.1 × 10^6^ greater than glucose.** As these calculations are based on total venom concentration, it is likely that the specific venom factors involved in sorbitol elevation are even more potent.

**Figure 3.**
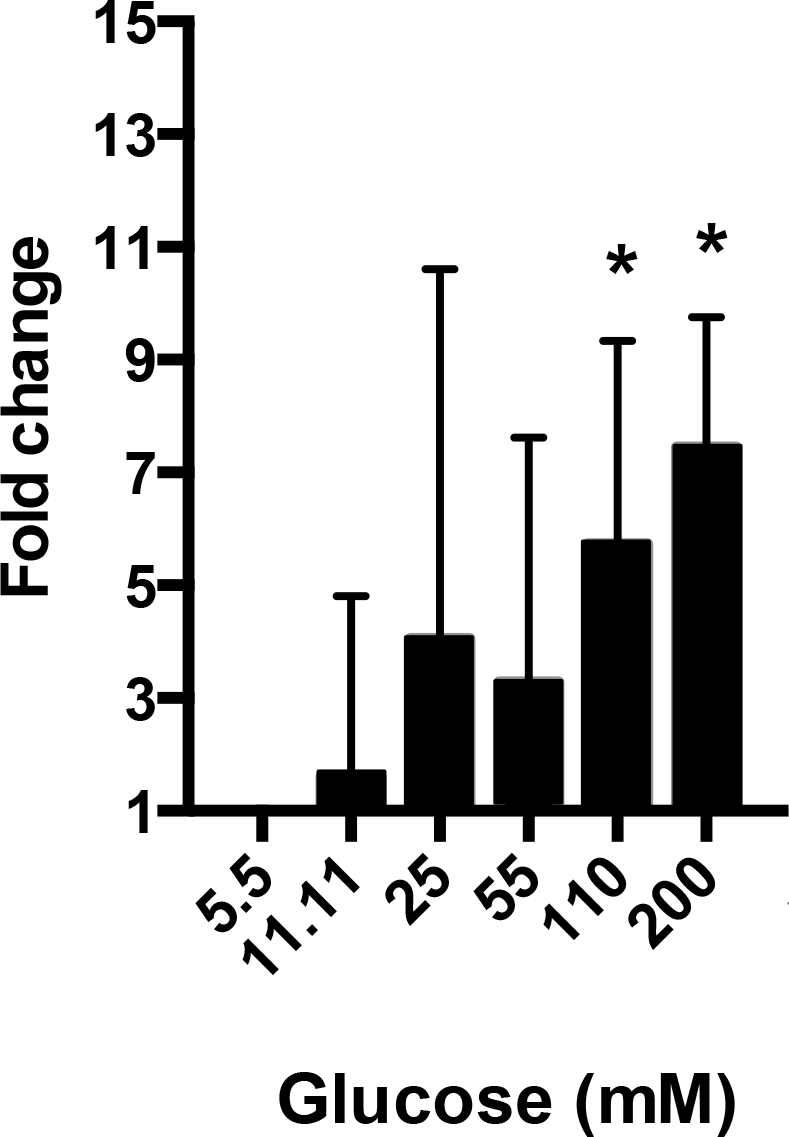
Moderate sorbitol elevation with high glucose treatment. High-dose glucose treatment elevates sorbitol level in HRMCs at 24 hours (n=4 samples per group) (*p<0.05). Fold changes are shown to the 5.5 mM and zero venom control.

### Low-dose venom alters expression of human metabolic genes in a pattern similar to that observed in the natural fly host

To initially survey genome wide gene expression changes in HRMCs exposed to different venom doses, single RNA-Seq replicates were employed for 1 hour, 4 hours and 24 hours after low-dose venom exposure (0, 1/128 VRE, 1/64 VRE, 1/16 VRE). At 1-hour post venom exposure, RNA-Seq showed no differentially expressed genes compared to control for any of the doses, suggesting that venom is not directly targeting preformed mRNA within the cells. After 4 hours a number of genes are differentially expressed (153 at 1/128 VRE, 95 at 1/64 VRE, and 182 at 1/16 VRE). In contrast, by 24 hours relatively fewer genes are differentially expressed at all venom doses compared to controls (67 at 1/128 VRE, 22 at 1/64 VRE, and 127 at 1/16 VRE).

Based on these results, we decided to focus attention on changes at 4 hours because they are most likely to reflect primary effects of venom that correspond to the observed sorbitol elevation at 24 hours post-exposure. Two additional biological replicates were sequenced for each of the sorbitol-elevating venom doses (1/128, 1/64, and 1/16 VRE) along with two additional experimental controls, for a total of 3 biological replicates per dose and 5 control samples. The number of significant venom responsive genes ranged from 137 (1/16 VRE) to 1605 (1/64 VRE) with the lowest dose (1/128) being intermediate (426) (Figure 4).

**Figure 4.**
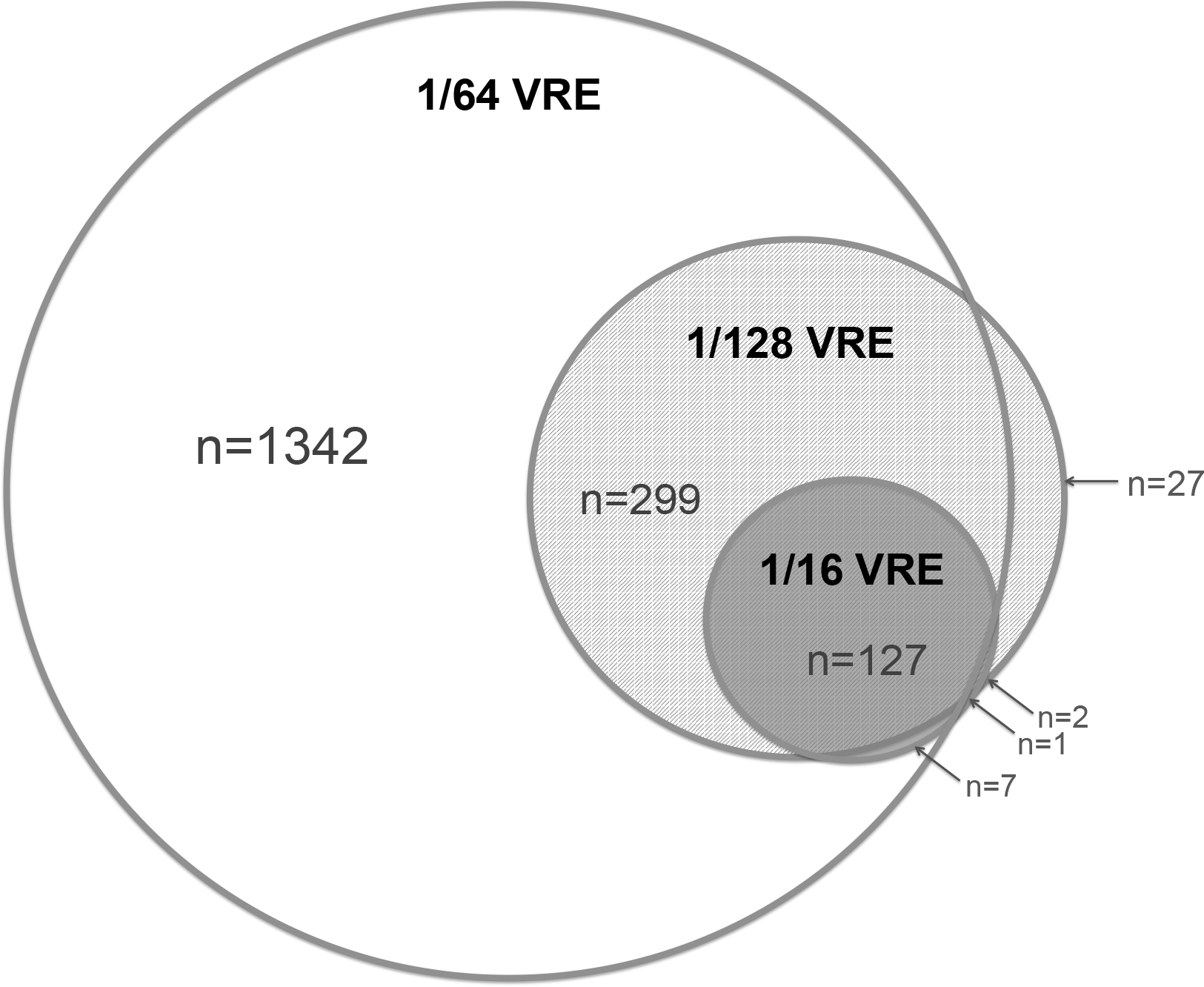
Relatively few genes (n=127) are consistently differentially expressed in response to sorbitol-elevating doses of *Nasonia venom*. Venn diagram of significantly differentially expressed genes (P<0.05, FDR<0.01) in human renal mesangial cells after 4 hours of exposure to low venom concentrations: 1/124, 1/64, 1/16 venom reservoir equivalents (VRE).

There is a set of 127 genes that were significantly differentially expressed across all 3 sorbitol-elevating doses (p<0.05, FDR<0.01, Figure 4); with 58 genes down-regulated and 69 up-regulated in response to venom treatment. The direction of change for each of these genes was the same across all three venom doses. This represents only 0.54% of transcripts detectable in HRMCs (127/23368 contigs). Gene ontologies of this overlapping set (n=127) cluster in a limited number of metabolic pathways, which includes up-regulation of sterol and small molecule metabolism, as well as modulation of chemical/stimulus response and cell signaling genes, and down-regulation of developmental and structural genes (Tables S1 & S2).

At 4 hour post treatment, up-regulated genes primarily function in metabolism, including alcohol (lanosterol synthase, insulin induced gene 1, acetyl-coa acetyltransferase 2, farnesyl diphosphate synthase), steroid hormones and cholesterol (nadp-dependent steroid dehydrogenase-like, 7-hydroxycholesterol reductase, 24-dehydroxycholesterol reductase, LDL receptor, and 2 sterol regulatory element binding transcription factors), and hydroxyl compounds (synuclein alpha, mevalonate decarboxylase). Down-regulated genes include those involved in extracellular matrix organization (matrix metallopeptidase 1, collagen type IV and V, chondroitin sulfate n-acetylgalactosaminyltransferase I, and gremlin), cell migration (fibroblast growth factor I, split homolog), chemotaxis (thrombospondin I, endothelin I, coagulation factor III), fatty acid transport (insulin receptor substrate 2), coagulation and regulation of wound healing (serpin peptidase inhibitor, heparin-binding egf-like growth factor, coagulation factor III).

A large number of DE genes are annotated as ‘Positive Regulation of Transcription from RNA Polymerase II Promoter’ (enrichment score 2, p=9.10E-04, E=3.67E-02). This includes both up-regulated (n=10) and down regulated (n=6) genes. The finding suggests a direct effect of *Nasonia* venom on transcriptional regulation in HRMCs and may explain the persistence of venom effects 24 hours post-exposure.

MT-RNR2-like 1 is the most significantly differentially expressed gene in this overlapping set. It is a mitochondrial-derived nuclear gene with sequence homology to Humanin 1. Humanin and its derivatives have been implicated in protection against oxidative stress mediated by superoxide dismutase (50, 51). Low dose venom causes between a −9.11 to −9.26 log2-fold decrease in expression of this gene at 4 hours, and this gene has the highest DE rank across all sorbitol-elevating doses of venom.

We also examined DE genes at 24 hours of exposure by pooling the three single-replicate, low-dose venom exposed samples (1/128, 1/64, and 1/16 VRE). At this time-point there were 47 differentially expressed genes; with 12 down-regulated and 35 up-regulated. Of these 26 were not differentially expressed at 4 hours. The remaiing 21 genes were differentially expressed at both 4 and 24 hours, with 18 showing the same direction of change (17 genes remain up-regulated and 1 gene remains down regulated) and 3 showing a significant reversal in expression (Table S3). All differentially expressed genes show a marked reduction in the magnitude of the fold change at 24 hours as compared to 4 hours. Gene ontology analysis of the full set of 47 genes again shows significant enrichment for alcohol and steroid metabolism, and hormone response (Table S3).

The broad pattern of change is similar to our previous findings of venom-induced expression effects in the natural fly host transcriptome, including alcohol, steroid metabolism, and hormone biosynthesis (9). The observation suggests strongly conserved venom functions between two species that are approximately 520 million years divergent from one another, the divergence time of deuterostomes and protostomes (52).

### Canonical sorbitol pathway genes AR and SD do not show expression changes, but AKR1C3 does

Both venom and glucose cause elevation of sorbitol at 24-hours. We therefore investigated sorbitol pathway gene expression changes that precede this elevation, for known and suspected genes involved in sorbitol metabolism (53). Neither the canonical AR (AKR1B1) nor the downstream enzyme in the sorbitol pathway, SD, showed significant up-regulation 4 hours after treatment with *Nasonia* venom. In contrast, sorbitol-elevating *Nasonia* venom doses (1/128, 1/64, 1/16 VRE = 1.2 × 10^−2^, 2.4 × 10^−2^, 9.6 × 10^−2^ μg/mL) cause significant up-regulation of the gene encoding an enzyme related to AR, AKR1C3 (log2-fold change and FDR of 1.28 at 1.95E-3, 1.20 at 9.44E-04, and 1.05 at 8.26E-03 respectively). These low venom dose responses were observed 4-hours after venom treatment, which is 20-hours prior to the detected sorbitol accumulation. Neither AR nor AKR1C3 are significantly differentially expressed at 24 hours in our single-replicate pilot experiment.

AKR1C3 belongs to the same protein family as AR but is not recognized as a sorbitol pathway gene. However, the gene has previously been identified amongst the top 50 genes differentially expressed *in vivo* in diabetic nephropathy (54). AKR1C3 is a member of a sub-family of aldo-keto reductase encoding genes, arranged in a syntenic group on human chromosome 10p15-p14, some of which are known to be transcriptionally co-regulated (55). Studies have focused on the function of AKR1C* sub-family enzymes in steroid hormone metabolism (56). Their role in sugar metabolism has not been investigated. These candidate sorbitol-pathway AKRs exhibit 48-49% protein sequence identity with the canonical sorbitol synthesizing enzyme AR (AKR1B1) (57).

Based on the RNASeq results, AKR1C3 is a potential candidate for involvement in sorbitol synthesis. To investigate the likelihood of enzyme interaction with sorbitol pathway substrates, we estimated relative binding affinity (K_m_) of candidate sorbitol pathway AKRs from humans, using *in silico* modeling of docking potential for glucose with NADPH cofactor bound. Docking potential is the predicted affinity of small molecules, in this case glucose, for a receptor of known or predicted 3-dimensional structure (48, 49). The more negative the estimate produced by iGemDocks, the greater the predicted binding affinity for the target substrate. Three AKR1C* sub-family enzymes (1C1, 1C2, 1C3) exhibited greater relative binding affinity for glucose than did the canonical sorbitol-synthesizing enzyme AR (AKR1B1) (Table 3). AKR1C3 had the highest predicted binding affinity for glucose (−70.46 relative to −65.04 for AR). AKR1C1, 1C2, 1C3 enzymes could therefore interact with elevated glucose more strongly than the low-affinity AR, which required high levels of substrate to catalyze the glucose to sorbitol conversion, both *in vitro* and *in vivo* (53). A role of these AKRs in glucose to sorbitol conversion in human insulin-insensitive cells has not been widely considered, but these results suggest that they warrant further investigation.

## Discussion

Here we present the first demonstration that parasitoid venom can affect a clinically important trait (sorbitol level) in human cells. The sorbitol pathway is thought to be a major contributor to pathogenesis of diabetic kidney disease (3, 26, 58). However, the effects of the sorbitol pathway on cellular dysregulation have been inextricable from other effects of hyperglycemia in previous *in vitro* and *in vivo* model systems (22, 37). *Nasonia* venom uncouples sorbitol pathway activation from hyperglycemia in human kidney cells *in vitro* and is a highly potent inducer of sorbitol elevation (2.1 × 10^6^ higher efficacy than glucose per unit weight). Venom also elevates sorbitol without elevating glucose in the natural fly host of *Nasonia* (1). Therefore, this demonstrates conservation of a parasitoid wasp venom phenotype in human cells with potential medical relevance. Direct interaction of venom proteins with sorbitol pathway enzyme(s) is a possible mechanism of action but would likely lead to immediate alteration of catalytic function. However, at 1 and 4 hours after venom exposure, we did not observe significant elevation of sorbitol levels, making this mechanism less likely. In contrast, venom treatment induces transcription of likely sorbitol pathway enzymes (AKRs) at 4 hours post exposure, which precedes intracellular sorbitol elevation observed at approximately 24-hours. These findings support a model of parasitoid wasp venom acting through a transcriptional mechanism to alter sorbitol metabolism in HRMCs under normoglycemic conditions.

*Nasonia* venom significantly elevates sorbitol in human renal cells in culture without altering levels of glucose or fructose. We examined gene expression changes occurring at 4 hours, 20 hours prior to the elevated sorbitol phenotype, to identify a putative mechanism. We focused our attention on the AKRs that have been implicated in sorbitol metabolism and/or diabetic nephropathy (59). One of these genes (AR) has traditionally been annotated as encoding the primary sorbitol-synthesizing enzyme. Although AR was not responsive to venom another related enzyme, AKR1C3, was significantly upregulated in response to low venom doses. AKR1C3 has been implicated in the pathophysiology of diabetic nephropathy - although no mechanism has been proposed (54).

Protein structural prediction and glucose docking reveal that AKR1C* enzymes have a relatively higher binding affinity for glucose than the canonical sorbitol synthesizing enzyme AR (AKR1B1). AKR1C3 may catalyze sorbitol synthesis *in vivo*, thereby contributing to sorbitol accumulation and glomerular dysfunction. This enzyme has been identified as one of fifty top genes up-regulated in the glomerulus of patients with diabetic nephropathy (54). AR has a low affinity for glucose and has been shown to catalyze sorbitol synthesis *in vitro* only when intracellular glucose levels are artificially elevated to levels much higher than would be experienced *in vivo* (60). Other AKRs may more efficiently catalyze sorbitol synthesis and therefore could also serve this function *in vivo* at physiologic intracellular glucose levels, comparable to those experienced by kidney cells during hyperglycemia (53, 61–63).

This study clearly reveals the potential of using *Nasonia* venom for studying sorbitol regulation in human cells, specifically to investigate how sorbitol elevation affects cellular mechanisms independent of elevated glucose. Next steps include identification of the causative venom component(s) and testing for molecular interactions both in the native host (*S. bullata*) and in human kidney cells. Additionally, identification of the molecular target of the venom would have important implications for (a) understanding sorbitol pathway regulation under altered metabolic conditions, (b) identifying the effect of sorbitol pathway activation independent of hyperglycemia, and (c) more effectively targeting therapeutics aimed at preventing sorbitol pathway activation and its various downstream effects on cellular function.

The genome-wide transcriptional analyses indicate that *Nasonia* venom alters early (4 hour) expression in a relatively small set of genes (127, or 0.54% of human genes that are enriched for metabolic categories, including sterol, alcohol, and hormone metabolism. Therefore, *Nasonia* venom constituents represent a potentially valuable source of metabolic effectors. More broadly, given the rapid turnover of venom repertoires in parasitoids (17, 64) and immense number of species (100,000 – 400,000), parasitoid venoms may represent a rich resource for discovery of biologics of medical and research relevance.

**Table 1.** Relative binding affinity for glucose of canonical and non-canonical AKR enzymes. *Homo sapiens* AKR1C* sub-family enzymes exhibit greater relative affinity for cyclical glucose than does the canonical sorbitol pathway enzyme Aldose Reductase (AR = AKR1B1). AKR1C3 has the highest predicted affinity for glucose (more negative value = greater binding affinity).

| Enzyme | Glucose | NADPH |
| --- | --- | --- |
| AKR1A1 | -64.10 | -118.40 |
| AKR1B1 (AR) | -65.04 | -124.77 |
| AKR1C1 | -69.58 | -103.59 |
| AKR1C2 | -66.42 | -65.054 |
| **AKR1C3** | **-70.46** | **-120.73** |
| AKR1C4 | -62.86 | -107.34 |

## Acknowledgements

We would like to thank A. Dolan, R. Edwards, M. He, E. Martinson, and Mrinalini for comments and discussion. This research was supported by the National Institutes of Health (F31DK108658 to ALS, RO1GM098667 to JHW), University of Rochester Drug Development Delivery Pilot Award to JHW and PJSH, and Helen Wisch Chair to JHW. Author contributions: ALS, PJSH, and JHW contributed to study design. ALS contributed to data collection. ALS, LAD, PJSH, and JHW contributed to data analysis and writing the paper.

